# Analysis of Serologic Cross-Reactivity Between Common Human Coronaviruses and SARS-CoV-2 Using Coronavirus Antigen Microarray

**DOI:** 10.1101/2020.03.24.006544

**Authors:** Saahir Khan, Rie Nakajima, Aarti Jain, Rafael Ramiro de Assis, Al Jasinskas, Joshua M. Obiero, Oluwasanmi Adenaiye, Sheldon Tai, Filbert Hong, Donald K. Milton, Huw Davies, Philip L. Felgner, Prometheus Study Group

## Abstract

The current practice for diagnosis of SARS-CoV-2 infection relies on PCR testing of nasopharyngeal or respiratory specimens in a symptomatic patient at high epidemiologic risk. This testing strategy likely underestimates the true prevalence of infection, creating the need for serologic methods to detect infections missed by the limited testing to date. Here, we describe the development of a coronavirus antigen microarray containing immunologically significant antigens from SARS-CoV-2, in addition to SARS-CoV, MERS-CoV, common human coronavirus strains, and other common respiratory viruses. A preliminary study of human sera collected prior to the SARS-CoV-2 pandemic demonstrates overall high IgG reactivity to common human coronaviruses and low IgG reactivity to epidemic coronaviruses including SARS-CoV-2, with some cross-reactivity of conserved antigenic domains including S2 domain of spike protein and nucleocapsid protein. This array can be used to answer outstanding questions regarding SARS-CoV-2 infection, including whether baseline serology for other coronaviruses impacts disease course, how the antibody response to infection develops over time, and what antigens would be optimal for vaccine development.

## Background

The 2019 novel coronavirus strain (SARS-CoV-2) originating in Wuhan, China has become a worldwide pandemic with significant morbidity and mortality estimates up to 2% of confirmed cases. The current case definition for confirmed COVID-19 due to SARS-CoV-2 infection relies on PCR-positive nasopharyngeal or respiratory specimens, with testing largely determined by presence of fever or respiratory symptoms in an individual at high epidemiologic risk. However, this case definition likely underestimates the prevalence of SARS-CoV-2 infection, as individuals who develop subclinical infection that does not produce fever or respiratory symptoms are unlikely to be tested, and testing by PCR of nasopharyngeal or respiratory specimens is unlikely to be 100% sensitive in detecting subclinical infection. Widespread testing within the United States is also severely limited by the lack of available testing kits and testing capacity limitations of available public and private laboratories. Therefore, the true prevalence of SARS-CoV-2 infection is currently unknown, and the sensitivity of PCR to detect infection is also unknown.

Serology can play an important role in defining both the prevalence of and sensitivity of PCR for SARS-CoV-2 infection, particularly for subclinical infection. This point is demonstrated by analogy with influenza virus, for which a meta-analysis of available literature measured the fraction of asymptomatic infections detected by PCR as approximately 16%, while the fraction of asymptomatic infections detected by seroconversion was measured as approximately 75% ^1^. The seroprevalence of common human coronaviruses is known to increase throughout childhood to near 100% by adolescence ^2^. Thus, any serologic methodology to estimate prevalence of SARS-CoV-2 needs to identify and rule out cross-reactivity with these common human coronavirus strains.

One challenge in applying serology to SARS-CoV-2 is that the choice of antigen and choice of assay is less well defined for coronavirus than more well studied viruses such as influenza. However, prior approaches to serologic detection of infection with emerging coronaviruses including SARS and MERS have focused on the S1 domain of the spike (S) glycoprotein and the nucleocapsid (N) protein, which are considered the immunodominant antigens for these viruses ^3^. In particular, the S1 domain is strain-specific, while the N protein shows cross-reactivity across strains.

The assay methodologies used for serologic detection of coronavirus infection include binding and neutralization assays. These methodologies have been shown to be well correlated ^4^. However, neutralization assays require viral culture, which must be performed in high-level biosafety containment units for emerging coronaviruses with high epidemic potential such as SARS-CoV-2. Conversely, binding assays such as ELISA can be readily performed with widely available reagents and equipment so are field deployable and suitable for point of care testing.

The protein microarray methodology has been widely used to simultaneously perform binding assays against hundreds of antigens printed onto a nitrocellulose-coated slide for detection of multiple antibody isotypes ^5^. This methodology was recently demonstrated for simultaneous measurement of IgG and IgA antibodies against over 250 antigens from diverse strains and subtypes of influenza ^6^. This methodology has previously been applied to detect antibodies against the S1 domains of SARS and MERS coronaviruses ^7^.

## Methodology

### Specimen Collection

The blood specimens used in this study were collected for a larger study where residents of a college resident community in an Eastern university were monitored prospectively to identify acute respiratory infection (ARI) cases using questionnaires and RT-qPCR, so as to characterize contagious phenotypes including social connections, built environment, and immunologic phenotypes^8^. From among de-identified blood specimens for which future research use authorization was obtained, five specimens that showed high IgG reactivity against human coronaviruses in the larger study were chosen for validation of the coronavirus antigen microarray.

### Antigen Microarray

The coronavirus antigen microarray used in this investigation includes 67 antigens across subtypes expressed in either baculovirus or HEK-293 cells (see Tables 1-3). These antigens were provided by Sino Biological Inc. (Wayne, PA) as either catalog products, or service products. The antigens were printed onto microarrays, probed with human sera, and analyzed as previously described (Figure 1)^6,9,10^.

**Table 1.** Content of coronavirus antigen microarray.

| Virus | Subtypes | Antigens | Replicates | Spots |
| --- | --- | --- | --- | --- |
| Coronavirus | HKU1, OC43, NL63, 229E | 12 | 4 | 48 |
|  | MERS | 9 | 4 | 36 |
|  | SARS | 5 | 4 | 20 |
|  | 2019-nCoV | 7 | 4 | 28 |
| Total |  | 33 |  | 132 |
| RSV | A, B | 8 | 4 | 32 |
| Metapneumovirus | A, B | 3 | 4 | 12 |
| Parainfluenza | 1, 3, 4 | 5 | 4 | 20 |
| Adenovirus | 3, 4, 7 | 6 | 4 | 24 |
| Influenza | H1N1, H3N2, H5N1, H7N9, B(Yam), B(Vic) | 12 | 4 | 48 |
| Total |  | 34 |  | 136 |

**Table 2.**
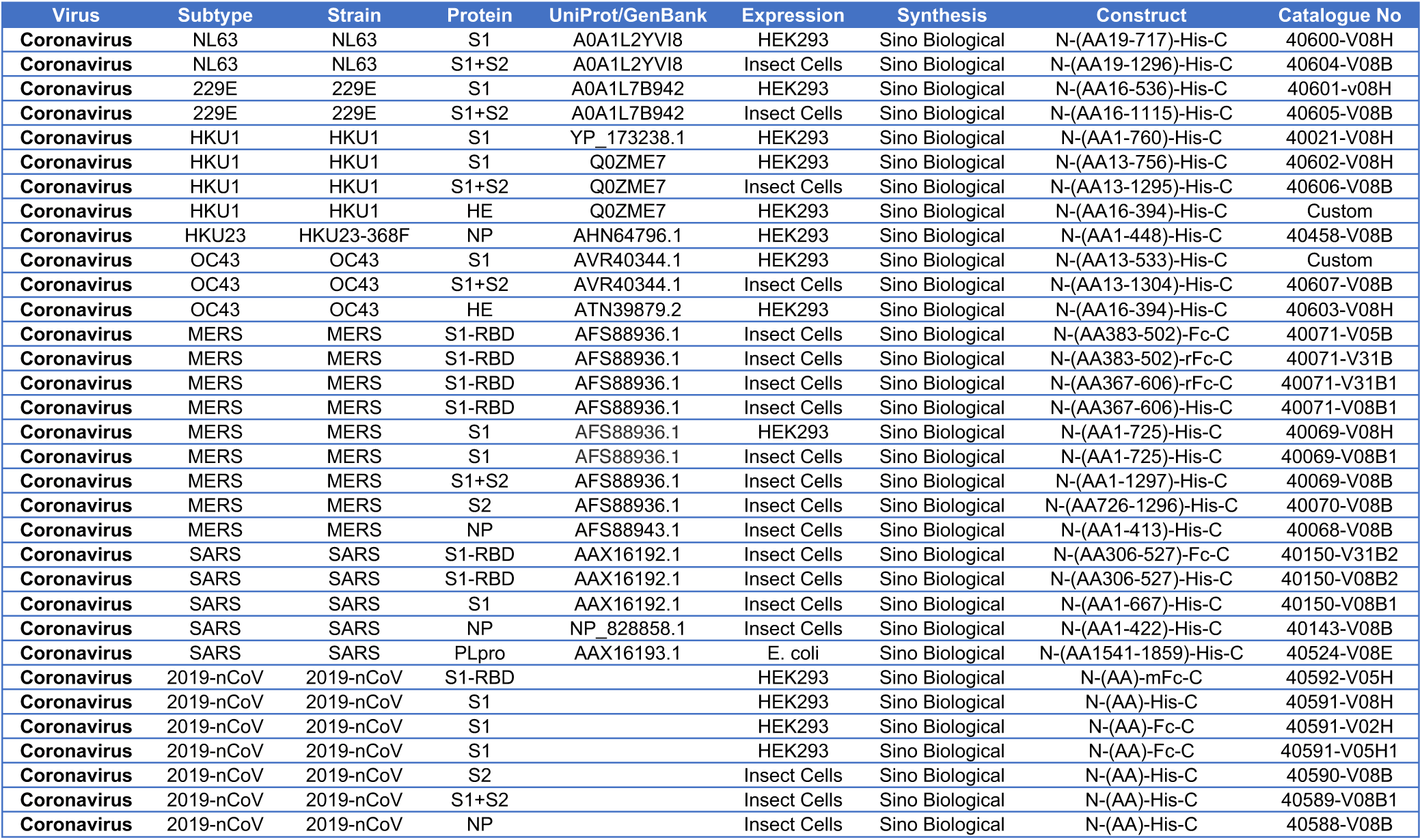
Coronavirus antigens on microarray.

**Table 3.** Non-coronavirus respiratory virus antigens on microarray.

| Virus | Subtype | Strain | Protein | UniProt/GenBank | Expression | Synthesis | Construct | Catalogue No |
| --- | --- | --- | --- | --- | --- | --- | --- | --- |
| RSV | A | LA2-94/2013 | F | A0A023RA53 | Insect Cells | Sino Biological | N-(AA1-526)-His-C | Custom |
| RSV | A | LA2-94/2013 | G | A0A076FRQ0 | HEK293 | Sino Biological | N-(AA64-321)-His-C | Custom |
| RSV | A | A2 | F |  | Insect Cells | Sino Biological | N-(AA1-529)-His-C | 11049-V08B |
| RSV | A | rsb1734 | G |  | HEK293 | Sino Biological | N-(AA66-297)-His-C | 11070-V08H |
| RSV | A | RSS-2 | F |  | Insect Cells | Sino Biological | N-(AA1-529)-His-C | 40037-V08B |
| RSV | B | TH-10526/2014 | F | K7WLI9 | Insect Cells | Sino Biological | N-(AA1-525)-His-C | Custom |
| RSV | B | TH-10526/2014 | G | A0A142MLK4 | HEK293 | Sino Biological | N-(AA64-310)-His-C | Custom |
| RSV | B | B1 | G |  | HEK293 | Sino Biological | N-(AA67-299)-His-C | 13029-V08H |
| hMPV | A | PER/CFI0320/2010/A | G |  | HEK293 | Sino Biological | 52N-228N-His | Custom |
| hMPV | B | PER/CFI0466/2010/B | G |  | HEK293 | Sino Biological | 52D-238S-His | Custom |
| hMPV | B | PER/CFI0320/2010/A | F |  | HEK293 | Sino Biological | 280D-490G-His | Custom |
| Parainfluenza | 1 | 12O3 | F | A0A1V0E1X5 | Insect Cells | Sino Biological | N-(AA22-497)-His-C | Custom |
| Parainfluenza | 1 | 12O3 | H | A0A1B2CW87 | Insect Cells | Sino Biological | N-His-(AA60-575)-C | Custom |
| Parainfluenza | 3 | USA/10991B/2010 | H | T1UD13 | Insect Cells | Sino Biological | N-His-(AA55-575)-C | Custom |
| Parainfluenza | 4 | hPIV-4b/10-H2/2016 | F | A0A1V0E1N6 | Insect Cells | Sino Biological | N-(AA22-486)-His-C | Custom |
| Parainfluenza | 4 | hPIV-4b/10-H2/2016 | H | A0A1V0E1N4 | Insect Cells | Sino Biological | N-His-(AA48-575)-C | Custom |
| Adenovirus | 3 | hAdV-3/45659 | Fiber | P04501 | E. coli | Sino Biological | N-His-[Prot]-C | Custom |
| Adenovirus | 3 | hAdV-3/45659 | Penton | Q2Y0H9 | Insect Cells | Sino Biological | N-His-[Prot]-C | Custom |
| Adenovirus | 4 | hAdV-4/28280 | Fiber | P36844 | Insect Cells | Sino Biological | N-[Prot]-His-C | Custom |
| Adenovirus | 4 | hAdV-4/28280 | Penton | Q2KSF3 | Insect Cells | Sino Biological | N-[Prot]-His-C | Custom |
| Adenovirus | 7 | Adeno7 10519 | Fiber | P15141 | Insect Cells | Sino Biological | N-His-[Prot]-C | Custom |
| Adenovirus | 7 | Adeno7 10519 | Penton | Q2KS58 | Insect Cells | Sino Biological | N-[Prot]-His-C | Custom |
| Influenza | H1N1 | A/Beijing/22808/2009 | HA1 | ADD64203.1 | HEK293 | Sino Biological | N-(AA1-344)-His-C | 40035-V08H1 |
| Influenza | H1N1 | A/Beijing/22808/2009 | HA1+HA2 | ADD64203.1 | HEK293 | Sino Biological | N-(AA1-529)-His-C | 40035-V08H |
| Influenza | H3N2 | A/Texas/50/2012 | HA1 | AGL07159.1 | HEK293 | Sino Biological | N-(AA1-345)-His-C | 40354-V08H1 |
| Influenza | H3N2 | A/Texas/50/2012 | HA1+HA2 | AGL07159.1 | Insect Cells | Sino Biological | N-(AA1-530)-His-C | 40354-V08B |
| Influenza | B | B/Malaysia/2506/2004 | HA1 | CO05957.1 | HEK293 | Sino Biological | N-(AA1-362)-His-C | 11716-V08H1 |
| Influenza | B | B/Malaysia/2506/2004 | HA1+HA2 | CO05957.1 | HEK293 | Sino Biological | N-(AA1-556)-His-C | 11716-V08H |
| Influenza | B | B/Phuket/3073/2013 | HA1 | EPI529345 | HEK293 | Sino Biological | N-(AA1-361)-His-C | 40498-V08H1 |
| Influenza | B | B/Phuket/3073/2013 | HA1+HA2 | EPI529345 | Insect Cells | Sino Biological | N-(AA1-547)-His-C | 40498-V08B |
| Influenza | H5N1 | A/Vietnam/1203/2004 | HA1 | AAW80717.1 | HEK293 | Sino Biological | (AA1-342)-mFcγ1-His | 10003-V06H1 |
| Influenza | H5N1 | A/Vietnam/1203/2004 | HA1+HA2 | AAW80717.1 | HEK293 | Sino Biological | (AA1-531)-mFcγ1-His | 10003-V06H3 |
| Influenza | H7N9 | A/Anhui/1/2013 | HA1 | AGJ51953.1 | HEK293 | Sino Biological | N-(AA1-338)-His-C | 40103-V08H1 |
| Influenza | H7N9 | A/Anhui/1/2013 | HA1+HA2 | AGJ51953.1 | HEK293 | Sino Biological | N-(AA1-524)-His-C | 40103-V08H |

**Figure 1.**
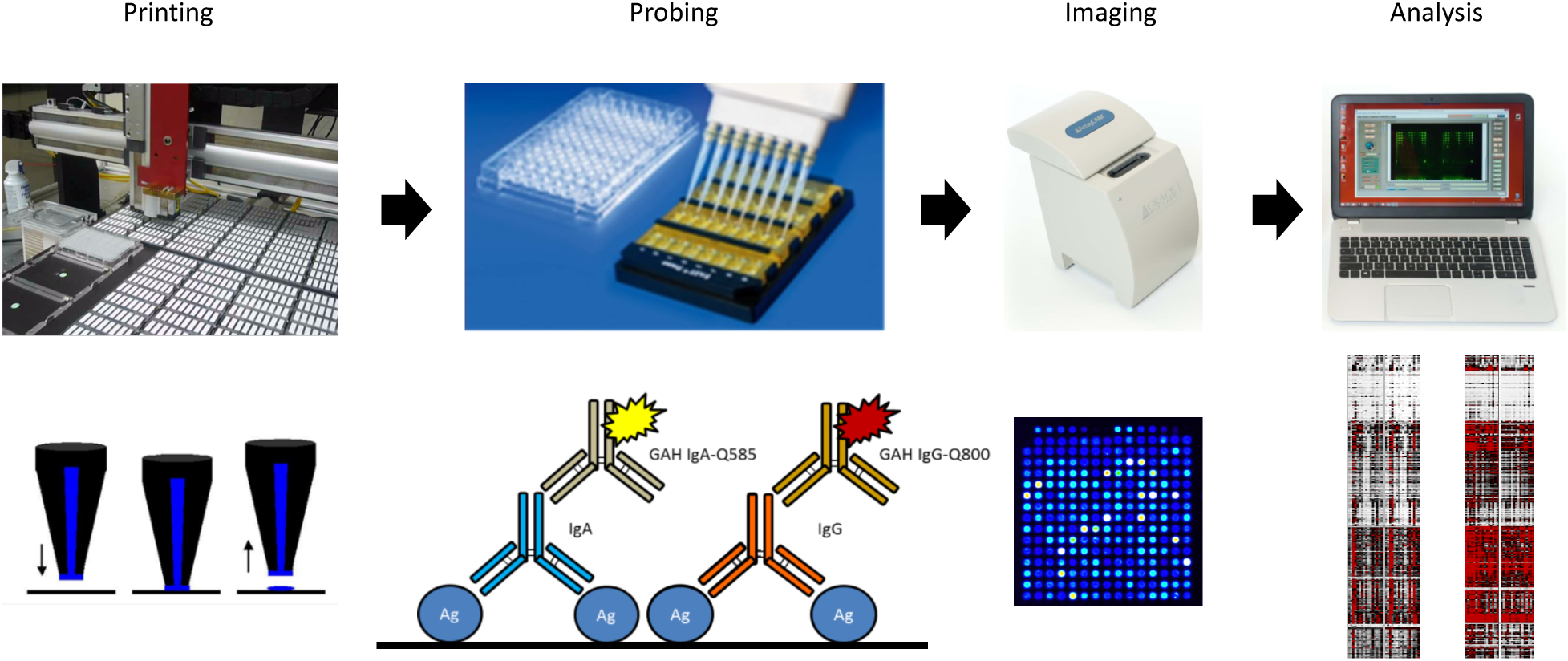
Schematic of antigen microarray printing, probing, imaging, and analysis. Reprinted with permission^6^.

Briefly, lyophilized antigens were reconstituted to a concentration of 0.1 mg/mL in phosphate-buffered saline (PBS) with 0.001% Tween-20 (T-PBS) and then printed onto nitrocellulose-coated slides from Grace Bio Labs (GBL, Bend, OR) using an OmniGrid 100 microarray printer (GeneMachines). The microarray slides were probed with human sera diluted 1:100 in 1x GVS Fast Blocking Buffer (Fischer Scientific) overnight at 4°C, washed with T-TBS buffer (20 mM Tris-HCl, 150 mM NaCl, 0.05% Tween-20 in ddH2O adjusted to pH 7.5 and filtered) 3 times for 5 minutes each, labeled with secondary antibodies to human IgG conjugated to quantum dot fluorophore for 2 hours at room temperature, and then washed with T-TBS 3 times for 5 minutes each and dried. The slides were imaged using ArrayCam imager (Grace Bio Labs, Bend, OR) to measure background-subtracted median spot fluorescence. Non-specific binding of secondary antibodies was subtracted using saline control. Mean fluorescence of the 4 replicate spots for each antigen was used for analysis.

### Statistical Analyses

Descriptive statistics were used to summarize the IgG reactivity as measured by mean fluorescence across antigen replicates. T-test or F-test were used to test for the mean differences in continuous variables across infection groups. All statistical analyses were conducted using R version 3.5.1 (R Foundation for Statistical Computing, Vienna, Austria).

## Results

Overall, the 5 sera tested on the coronavirus antigen microarray all showed high IgG seroreactivity to antigens from common human coronaviruses and other respiratory viruses with known seasonal circulation versus low IgG seroreactivity to antigens from epidemic viruses that were not circulating at time of collection (Figure 2). Specifically, 4 of the 5 sera showed high IgG seroreactivity across the 4 common human coronaviruses, while all of the sera showed low IgG seroreactivity to SARS-CoV-2, SARS-CoV, and MERS-CoV. All 5 sera showed high IgG seroreactivity to RSV and parainfluenza viruses, while 3 of the 5 sera showed high IgG seroreactivity to adenoviruses. For influenza, all 5 sera showed high IgG seroreactivity to H1N1 and H3N2 influenza A and influenza B strains but low IgG seroreactivity to H5N1 and H7N9 influenza A strains.

**Figure 2.**
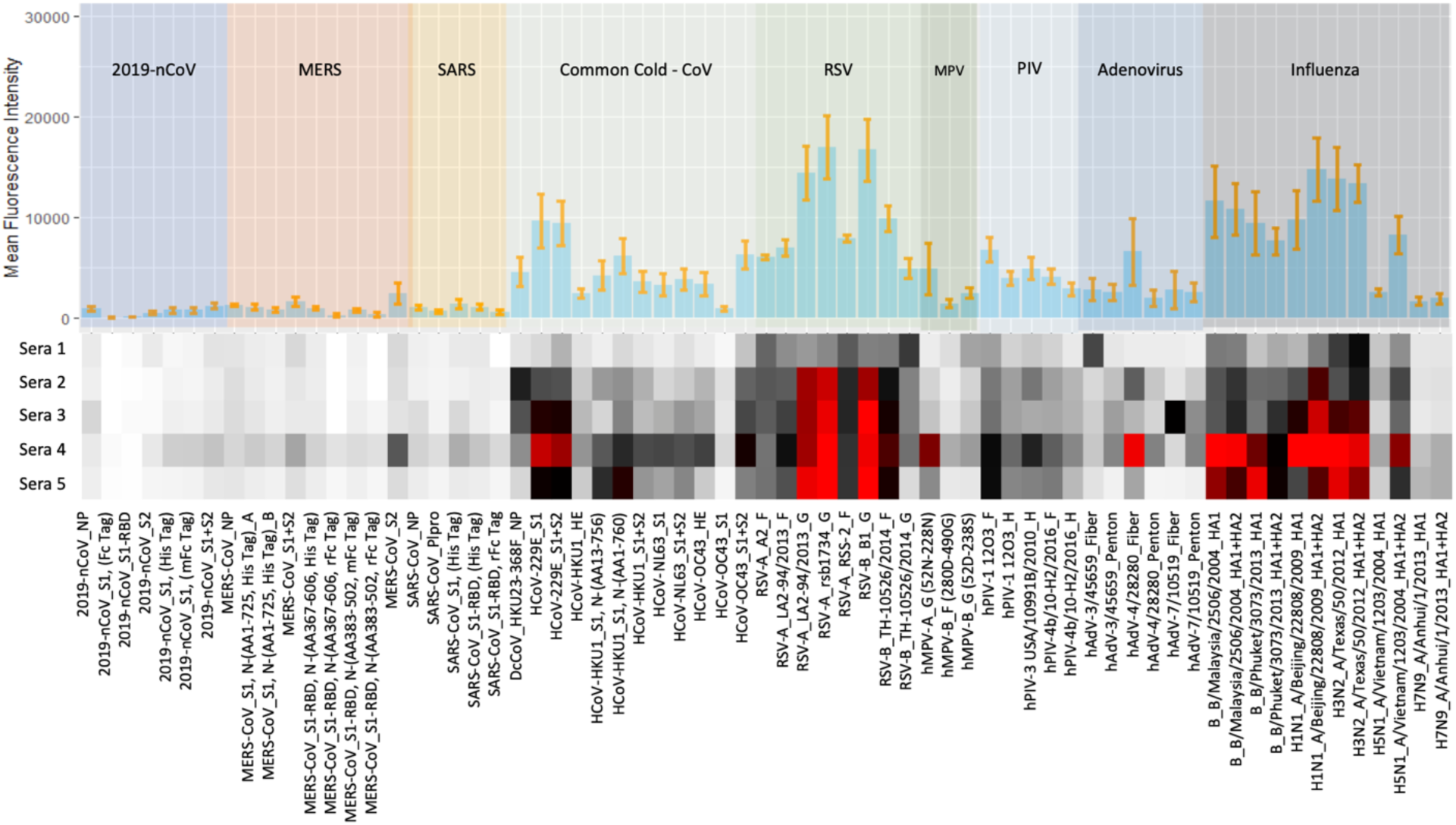
IgG seroreactivity as measured by mean fluorescence intensity of 5 serum specimens from naïve population on coronavirus antigen microarray.

With respect to specific antigens, the S1 domain of the spike protein including the receptor-binding domain (RBD) demonstrates very low cross-reactivity between epidemic coronaviruses and common human coronaviruses, whereas the S2 domain of the spike protein and the nucleocapsid protein (NP) show low-level cross-reactivity between these coronavirus subtypes. Similarly, the head domain of influenza hemagglutinin (HA1) is not cross-reactive between seasonal and avian influenza strains, whereas the stalk domain (HA2) is cross-reactive between influenza virus subgroups, as seen between H1N1 and H5N1 influenza viruses.

## Discussion

This pilot study yields several insights into cross-reactivity of common human coronavirus antibodies for SARS-CoV-2 antigens. The antibodies to the S1 and RBD domains of spike protein are highly subtype-specific, consistent with the high variability in these sequences between different human coronaviruses. Conversely, the S2 domain of spike protein and NP protein are more cross-reactive, consistent with these sequences being highly conserved across coronaviruses. SARS-CoV-2 has caused a worldwide pandemic despite likely pre-existing cross-reactive antibodies to S2 domain and NP protein in most people, indicating that these antibodies are likely not protective, whereas antibodies to S1 and RBD domains are more likely to be protective. This observation favors a vaccination strategy based on S1 or RBD domains of spike protein over a vaccination strategy that also includes S2 domain or NP protein. In addition, S1 and RBD domains are more likely to generate subtype-specific serologic tests for population surveillance studies.

In addition, a key unexplained finding during the SARS-CoV-2 epidemic has been the low incidence of infection in children aged 15 and younger. This observation generates two related hypotheses: adults may have pre-existing antibodies against antigenically distinct coronaviruses that produce an ineffective humoral response to SARS-CoV-2 infection (antibody-dependent enhancement as demonstrated for dengue virus), or children younger than 15 may have initially encountered a coronavirus that is more closely related to SARS-CoV-2 so are more protected against this infection (immunologic imprinting or original antigenic sin as demonstrated for influenza virus). Both of these hypotheses would be informed by comparing the level of cross-reactive coronavirus antibodies in pediatric and adult cohorts and correlating these antibodies with incidence of severe disease.

## Conclusions

A coronavirus antigen microarray has been constructed with antigens from epidemic coronaviruses including SARS-CoV-2 and common human coronaviruses, in addition to other common respiratory viruses. A pilot study of 5 naïve human sera shows high IgG seroreactivity to common human coronaviruses but low IgG seroreactivity to SARS-CoV-2, with some cross-reactivity seen for S2 domain of spike protein and nucleocapsid protein. Further studies are needed including with SARS-CoV-2 convalescent sera to fully realize the potential of this novel methodology to characterize the seroprevalence of SARS-CoV-2 and the impact of pre-existing cross-reactive antibodies on the disease course.

## Acknowledgements

Saahir Khan is supported by the National Center for Research Resources and the National Center for Advancing Translational Sciences, National Institutes of Health, through Grant KL2 TR001416. The content is solely the responsibility of the authors and does not necessarily represent the official views of the NIH. Prometheus-UMD was sponsored by the Defense Advanced Research Projects Agency (DARPA) BTO under the auspices of Col. Matthew Hepburn through agreements N66001-17-2-4023 and N66001-18-2-4015 (PI: Milton). This study was funded in part by the Defense Threat Reduction Agency via grants HDTRA1-18-1-0036 (PI: Davies) and HDTRA1-18-1-0035 (PI: Felgner). The findings and conclusions in this report are those of the authors and do not necessarily represent the official position or policy of the funding agencies and no official endorsements should be inferred.

